# Dose Escalation in Pentylenetetrazol Kindling Detects Differences in Chronic Seizure Susceptibility

**DOI:** 10.1101/2025.01.29.635374

**Authors:** Mitchell B. Moyer, Jenna Langbein, Orest Tsymbalyuk, Darrian McAfee, Chixiang Chen, Muznabanu Bachani, Volodymyr Gerzanich, J. Marc Simard, Alexander Ksendzovsky

## Abstract

**Objective:** Pentylenetetrazol (PTZ) kindling is a widely used model for inducing epileptogenesis and evaluating long-term seizure susceptibility differences among animals. This model is typically performed by chronic, repetitive exposures to a constant subconvulsive PTZ dose. However, the effectiveness of the commonly used subconvulsive dose (35mg/kg) varies among different animal groups and experimental conditions due to factors such as species, age, sex, and genetic background. The objective of this study was to characterize a novel model of kindling, the PTZ Dose Escalation (PTZ-DE) model, which assesses chronic seizure threshold with enhanced sensitivity by empirically determining the minimally effective dose to induce PTZ kindling for specific experimental conditions.

**Methods:** This study investigated the efficacy and validity of the PTZ-DE model by comparing its performance to the standard PTZ kindling approach across a series of conditions. First, the ability of the PTZ-DE model to produce the gradual increase in chronic seizure severity response characteristic of PTZ kindling was compared to the standard model across animal background characteristics (strain, sex). Next, the validity of this model was investigated by determining if the PTZ-DE model could replicate similar changes in chronic seizure susceptibility previously published using the standard approach after traumatic brain injury (TBI). Lastly, the PTZ-DE model’s efficacy to detect seizure differences was measured in a condition (glyburide treatment) in which alterations to chronic seizure susceptibility were not detected with standard kindling.

**Results:** This study found that the PTZ-DE model corrects for background differences in PTZ susceptibility, replicates known differences in chronic seizure thresholds after TBI, and identifies new alterations in seizure threshold not detected with traditional kindling methods.

**Significance:** The PTZ-DE model may prove to be a superior tool to standard PTZ kindling for discovering new pathological mechanisms of epileptogenesis and for developing targeted therapies for chronic seizure management, as evidenced by its ability to detect subtle differences in seizure susceptibility across various experimental conditions.

## 1. Introduction

Pentylenetetrazol (PTZ) kindling serves as the gold standard animal model for inducing epileptogenesis, uniquely increasing chronic seizure susceptibility without relying on status epilepticus (*1, 2*). This model is invaluable for investigating how chronic seizures impact the brain or examining how factors such as injury, pharmacological interventions, or genetic modifications affect chronic seizure susceptibility (*1, 2*). The standard PTZ kindling model is performed by intraperitoneal (IP) injection a fixed dose of 35mg/kg assumed to be a subconvulsive, every second day for twenty total days (*3*). Repetitive exposure to this initially subconvulsive dose eventually leads to a chronically lowered seizure threshold, which results in severe seizure responses to the same low-dose PTZ exposure (*3*). However, the subconvulsive nature of the 35mg/kg dose may vary across different animal groups or experimental conditions, as an animal’s response to PTZ is influenced by variables including species, age, sex, and genetic background (*4-7*). Therefore, to enhance the PTZ kindling model’s sensitivity in detecting differences in chronic seizure susceptibility among experimental groups, it may be important to empirically determine the minimal effective dose to induce PTZ kindling within the context of each experiment, rather than relying on predetermined assumptions.

In the present study, we introduce a novel PTZ kindling model, referred to as the PTZ Dose Escalation (PTZ-DE) model. We evaluate its efficacy across several applications where nuanced responses to PTZ are required (genetic, model, pharmacological and sex differences), aiming to replicate known chronic seizure susceptibility differences observed with traditional PTZ kindling and to identify previously undetected differences with standard PTZ kindling. We developed the PTZ-DE model by incorporating a dose escalation approach to assess chronic seizure susceptibility. Inspired by methods used in acute seizure threshold testing—which gradually increase the PTZ dose until an acute seizure occurs—this model adapts the strategy to evaluate susceptibility to chronic seizures (*8*). The gradual escalation of the PTZ dose enables a more precise determination of the minimally effective dose for specific experimental conditions. Consequently, the PTZ-DE model enhances the sensitivity of chronic seizure susceptibility assessment compared to standard PTZ kindling. By using an empirical approach to dose determination, it enables the detection of subtle differences in seizure susceptibility that might be overlooked by conventional methods and may uncover novel therapeutic targets for seizure management in epilepsy.

## 2. Materials and Methods

### 2.1. Ethics statement

Animal experiments comply with the ARRIVE guidelines and were performed under a protocol approved by the Institutional Animal Care and Use Committee of the University of Maryland School of Medicine, and in accordance with the relevant guidelines and regulations as stipulated in the United States National Institutes of Health Guide for the Care and Use of Laboratory Animals.

### 2.2. Subjects

C57/Bl6N mice were males and females aged 6-8 weeks obtained from Charles River Laboratories, whereas C57/Bl6N+129/SvJ mice were male and female mice aged 10-16 weeks and obtained from our breeding colony. For sex comparison experiments, both males and females were C57/Bl6N mice aged 6-8 weeks. For traumatic brain injury (TBI) experiments, C57/Bl6N mice were obtained from Envigo and were 8–12-week-old males. Glyburide (GLYB) and vehicle (VEH) treated C57/Bl6N mice were male and female aged 6-8 weeks old obtained from Charles River Laboratories.

### 2.3. Racine Scoring

Seizure severity scoring was monitored for 30 minutes following all PTZ injections as previously described using the Racine scoring scale (*9, 10*). In brief, the following classifications were used: 0. No seizure response, 1. Behavioral arrest or slowing, 2. Head nodding associated with facial clonus, 3. Partial limb clonus with or without tail stiffening, 4. Clonic seizure with rearing and loss of posture, 5. Generalized tonic-clonic seizure with wild running and/or jumping, 6. Death. After death, mice were assigned a Racine score of 6 for each day for the remainder of the experiment to prevent sampling bias and maintain statistical power between groups for the entire experiment.

### 2.4. PTZ Kindling (Standard)

Standard PTZ kindling was induced as previously described (*3*). In brief, PTZ (35mg/kg) was administered to mice every second day for a total of twenty days, i.e. 10 total PTZ administrations.

### 2.5. PTZ-DE Model

To evaluate differences in chronic seizure susceptibility, we modified the standard PTZ kindling model by initiating with a lower starting dose of 15mg/kg and incrementally increasing this dose by 5mg/kg after every three administrations. This escalation continued until at least one animal from any group exhibited Racine stage 4 or higher seizure response. Following this, the dose that elicited a Racine 4+ response was consistently used for subsequent PTZ administrations until group responses stabilized near a “fully kindled” state (Racine 4+) for at least 3 doses. PTZ administrations were still conducted every other day, as in the standard protocol, but with a variable total number of doses administered, typically ranging from 20 to 25.

### 2.6. Drug Administration

PTZ (Millipore-Sigma, P6500) was reconstituted in 1X PBS to a stock concentration of 10mg/ml and syringe filtered for sterility, after which PTZ was administered to mice via IP injection. For GLYB experiments, 10mg of GLYB (Millipore-Sigma, G2539) was weighed and reconstituted in 1ml of 100% corn oil (VEH) and diluted to 0.3mg/ml, after which 100μL of glyburide (final dose of 30μg/mouse) was administered IP daily concurrently with PTZ treatments. On PTZ administration days, GLYB was given 15 minutes prior to PTZ injection.

### 2.7. Controlled Cortical Impact (CCI) Induction

Mice underwent a controlled cortical impact (CCI) to model moderate to severe traumatic brain injury (TBI) (*11*). Briefly, mice were anesthetized with 4% isoflurane in oxygen and placed in a stereotactic frame (Stoelting Co., 51615). A left scalp incision was made. A dental drill was used to make a 5-mm craniotomy over the left parietal cortex, and the bone flap was removed. CCI was performed with an electromagnetic CCI impactor (Leica Biosystems, 39463920) with a flat tipped 3-mm diameter impounder on the left parietal cortex (velocity=1 m/s, depth = 2.5 mm, dwell time = 200 ms). Mice recovered in a temperature-controlled chamber before returning to their cage. Sham control mice underwent the identical procedure up until removal of bone; however, no impaction was induced. PTZ-DE was initiated 7 days after CCI.

### 2.8. Statistical analysis

Data are presented as mean ± SEM unless noted otherwise. Non-linear regression analyses using a logistic growth model with an extra sum-of-squares F test were used to assess for significant differences between PTZ kindling severity curves. Logistic growth was chosen as the most appropriate model for all experiments (excluding Kaplan-Meier analyses), as the sigmoidal curve inferred when using a logistic growth model best represents the expected phases of PTZ kindling, which include the initial minimal responsive to PTZ due to the subconvulsive nature of the dose, followed by the growth phase in which mice begin exhibiting a seizure response to PTZ, and finally the plateau phase in which mice have fully kindled (*3*). To assess which initial responses (Y_0_) were closest to zero, Y_0_ values were compared between curves by applying Akaike’s Information Criterion (AICc) to evaluate two candidate models: one with the setting 0<y0>0.2 and one with Y_0_≥0.2. The model with the smaller AICc value is statistically preferred. Growth rate constants and maximal values (Ymax) were assessed by logistic growth model with an extra sum-of-squares F test for figures 2 and 3 respectively. Additionally, R^2^ analysis was used to assess how well the logistic growth model fits the data. from Log-rank tests were performed to assess statistical differences between Kaplan-Meier curves. A complete summary of statistical analyses and their resulting descriptive statistics can be found in Supplemental Table 1. Analyses were performed with GraphPad Prism 9.3.1. P<0.05 was deemed to be statistically significant.

## 3. Results

### 3.1. PTZ-DE Better Models PTZ Kindling Through Empirical Subconvulsive Dose Determination

To determine whether the PTZ-DE model can identify alterations in seizure susceptibility to PTZ caused by background differences among mice, we directly compared the responses of mice subjected to PTZ-DE with those treated using the standard PTZ protocol. In an initial assessment of chronic seizure susceptibility using the standard PTZ kindling model, C57/Bl6N mice were compared to C57/Bl6N+129/SvJ mice, as 129/SvJ strain mice are commonly used for breeding genetic knockout mouse lines (*12, 13*). Compared to C57/Bl6N mice (Y_0_=0.616), mice bred on a C57/Bl6N+129/SvJ background exhibited significantly elevated initial seizure responses (Y_0_=2.076, p<0.001; F=8.964) to the first 35mg/kg dose of PTZ, with sustained high Racine scores throughout the experiment (Figure 1A). This unexpected response suggested that the conventional 35mg/kg dose may not accurately capture the gradual increase in seizure severity that PTZ kindling aims to model, where the initial response (Y_0_) should be close to 0 across all conditions and progressively elevate over the course of the kindling process. Consequently, we used the PTZ-DE model to determine and compare the true minimally effective dose for these genetic backgrounds (Figure 1B). To ensure the minimal dose was captured, we started with 15mg/kg—significantly lower than typically reported in PTZ kindling literature-and incrementally increased the dose after every three administrations until a Racine 4+ seizure response was observed in at least one mouse from either group. After the first Racine 4+ seizure, this empirically determined minimally effective dose was maintained for the rest of the experiment. A Racine score of 4 in at least one animal was used to define the appropriate PTZ dose because this represents the point where at least one animal reached a “kindled” state characterized by development of a full tonic-clonic seizure rather than a single convulsive episode. Once the first animal exhibits a kindled response to the empirically determined dose, it can be reasonably expected that the other animals with a given background will also appropriately respond to kindling, thus that dose is continued for the remainder of the experiment. Both C57/Bl6N and C57/Bl6N+129/SvJ mice appropriately displayed minimal seizure responses (Racine 0-1) (C57/Bl6N: Y_0_=0.0029; C57/Bl6N+SvJ: Y_0_=0.1681) at 15mg/kg and 20mg/kg PTZ, but C57/Bl6N+129/SvJ mice began to show increasing Racine scores from 25mg/kg onwards, whereas C57/Bl6N mice only began exhibiting Racine 4+ seizure responses after reaching 30mg/kg. These observations led to the conclusion that the most appropriate PTZ dose for kindling C57/Bl6N+129/SvJ mice is 25mg/kg, which is significantly lower than the standard 35mg/kg dose, and 30mg/kg for C57/Bl6N mice. This finding explains the initially severe response of C57/Bl6N+129/SvJ mice compared to C57/Bl6N mice under the standard model.

**Figure 1:**
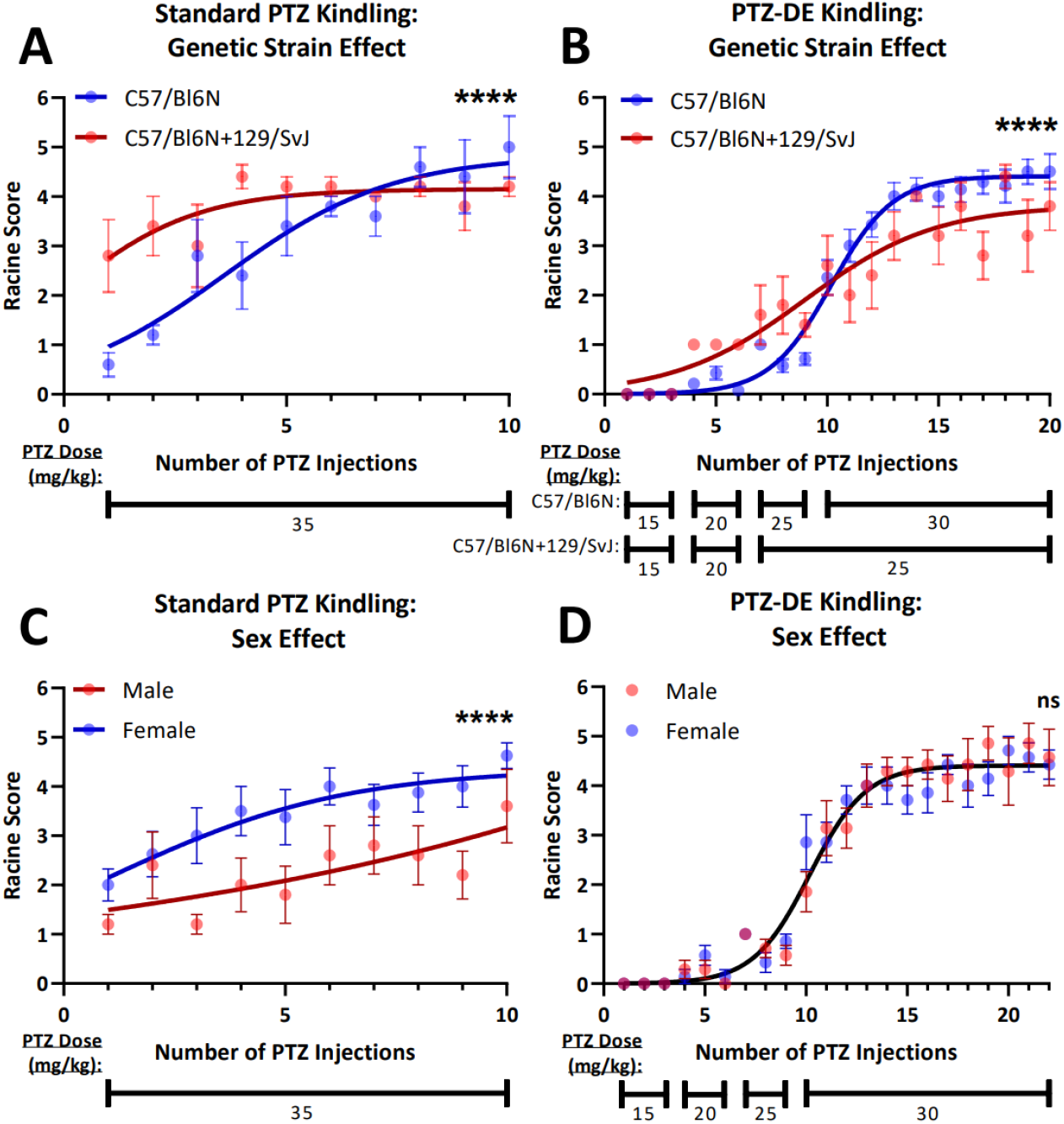
PTZ-DE Model Determines the Minimally Effective PTZ Dose to Correct Variabilities in PTZ kindling. A) Mice bred on a C57Bl/6N+129/SvJ background (n=5) exhibit elevated initial response compared to C57/Bl6N mice (n=5) in standard PTZ kindling. B) The PTZ-DE model empirically determines the true minimally effective dose to recreate gradually increasing seizure response to PTZ over the course of induction. C57Bl/6N mice (n=14) optimally kindle at 30mg/kg, whereas C57Bl/6N+129/SvJ mice (n=5) kindle at 25mg/kg. C) C57/Bl6N female mice (n=8) demonstrate heightened response to standard PTZ kindling compared to males (n=5). D) When kindled using PTZ-DE up to 30mg/kg, no difference in male (n=7) and female (n=7) responses is seen. ns: not significant, ****p<0.0001.

In addition to identifying the true minimally effective dose for kindling, the PTZ-DE model more accurately replicated the characteristic pattern of PTZ kindling, marked by an initial minimal response, a gradual increase in seizure severity and eventual plateau (Supplemental Table 1). When fitted to the logistic growth model, which represents the characteristic response of PTZ kindling, PTZ-DE curves demonstrated improved measures of fit compared to the standard model (PTZ-DE: R^2^: C57/Bl6N: 0.8262, C57/Bl6N+129/SvJ: 0.6366; standard: R^2^: C57/Bl6N: 0.5826, C57/Bl6N+SvJ: 0.1744). Furthermore, the initial responses of both strains were appropriately near zero in the PTZ-DE models (C57/Bl6N: Y_0_=0.0029; C57/Bl6N+129/SvJ: Y_0_=0.1681, AICc analysis preferred the 0<Y_0_<0.2 setting models for both strains), whereas mice undergoing standard kindling exhibited elevated initial responses (C57/Bl6N: Y_0_=0.616; C57/Bl6N+129SvJ: Y_0_=2.076, AICc analysis preferred the Y_0_≥0.2 setting models for both strains). Together, these data suggest that the PTZ-DE model more accurately replicates the expected PTZ kindling response, especially during the early phase of kindling, compared to the standard model.

Although the PTZ-DE model reduced the group variability in PTZ response due to genetic strain differences, and more closely produced the expected gradual increase in seizure severity than standard PTZ kindling, the PTZ-DE kindling curves exhibited by C57/Bl6N and C57/Bl6N+129/SvJ mice were still significantly different (p<0.001, F=14.47) due to the differing minimally effective doses by each genetic strain (Figure 1B). To that end, we tested whether the PTZ-DE model could completely normalize differences in another background characteristic - sex -seen in standard PTZ kindling (Figure 1C). Male C57/Bl6N mice undergoing standard PTZ kindling demonstrated a significantly decreased (p<0.001, F=10.86) kindling response compared to female C57/Bl6N mice. Additionally, while female mice achieved full kindled status as a group (Ymax: 4.368) during standard kindling, male mice failed to fully kindle within the timeframe of the kindling protocol. This is further supported by the infinitely high (unstable) Ymax value determined by the logistic growth curve, indicating that the male group response had not yet plateaued (Supplemental Table 1). Using the PTZ-DE model, male and female mice exhibited comparable responses (p=0.235, F=1.427), characterized by minimal seizure phenotypes (Racine 0-1) until a PTZ dose of 30mg/kg, after which both sexes began to develop severe seizure responses at similar rates (Figure 1D). Both groups ultimately achieved fully kindled status (Ymax: Male: 4.544; Female: 4.265). Fu rthermore, as observed with genetic strains, PTZ-DE curves for both sexes demonstrated improved fit with the logistic growth model (R^2^: Male: 0.8158; Female: 0.8436) and a lower initial starting point (Y_0_: Male: 0.002685; Female: 0.002089, AICc analysis preferred the 0<Y_0_<0.2 setting models for both sexes) compared to the standard model (R^2^: Male: 0.1670, Female: 0.2508; Y_0_: Male: 1.374; Female: 1.744, AICc analysis preferred the Y_0_≥0.2 setting models for both sexes) (Supplemental Table 1). These findings suggest that the PTZ-DE model can fully normalize some differences in PTZ kindling responses due to animal background characteristics and further supports the notion that PTZ-DE induces a more accurate kindling response.

### 3.2. PTZ-DE Detects Chronic Seizure Threshold Differences Previously Identified by Standard PTZ Kindling

To validate the efficacy of the PTZ-DE model in detecting differences in chronic seizure susceptibility, we used dose escalation in a scenario where such differences in chronic seizure responses are expected and have been previously described in standard kindling: TBI induced by a CCI model. Previous studies using standard PTZ kindling methods had demonstrated reduced seizure thresholds in CCI mice compared to sham controls (*14-16*). Using the PTZ-DE kindling approach, we assessed whether it could similarly detect alterations in seizure responses induced by CCI, thereby testing the validity of the PTZ-DE model. We observed that mice in both groups showed minimal responses to PTZ doses of 15mg/kg or 20mg/kg; however, at 25mg/kg, CCI mice exhibited significantly greater seizure responses (p<0.001, F=16.29), characterized by a substantially faster elevation in Racine scores (Growth Rate Constant (GRC): Sham: 0.1462, TBI: 0.5483; p=0.001, F=10.80, extra sum-of-squares F test) (Figure 2A, Supplemental Table 1). The minimal response observed until the 25mg/kg dose indicates that this was the appropriate minimally effective PTZ dose for kindling in our TBI model. Additionally, the severe Racine response in CCI mice was accompanied by a higher mortality rate due to seizures (p=0.0297) compared to sham control mice (Figure 2B). Together, these results obtained using the PTZ-DE model align with prior findings from models using the standard PTZ model (*14-16*), corroborating both the efficacy of PTZ-DE model in detecting differences in chronic seizure susceptibility and previous evidence that TBI lowers the chronic seizure threshold.

**Figure 2:**
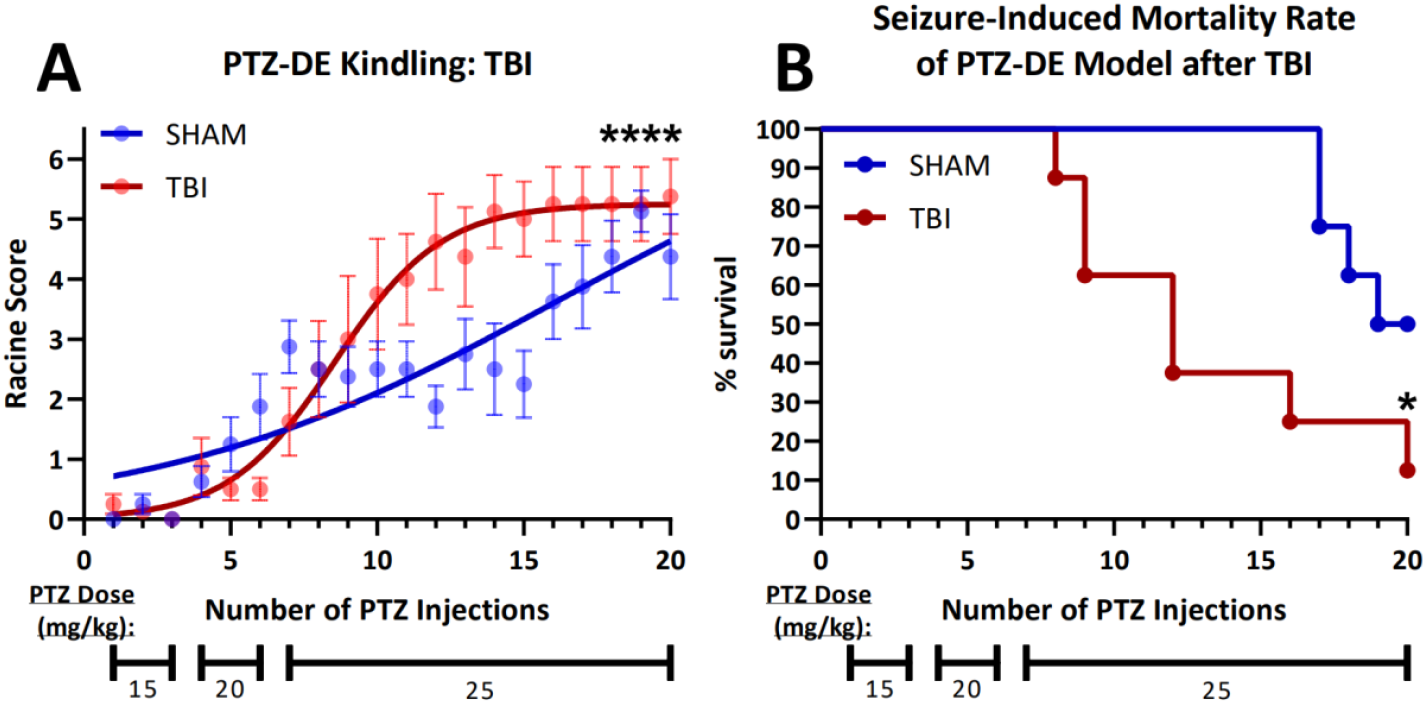
The PTZ-DE Model Replicates Elevated PTZ Kindling Response After TBI. A) TBI induced mice (n=8) develop more severe seizure responses with fewer PTZ administrations in the PTZ-DE model compared to sham controls (n=8). B) Kaplan-Meier analysis demonstrating increased mortality due to PTZ induced seizures in TBI mice (n=8) compared to sham controls (n=8) when assessed via PTZ-DE. *p<0.05, ****p<0.0001.

### 3.3 PTZ-DE Enhances Sensitivity for Detecting Chronic Seizure Susceptibility Differences Compared to Standard PTZ Kindling

To directly compare the PTZ-DE model’s ability to detect nuanced differences in seizure susceptibility due to pharmacological interventions to the standard kindling approach, we used both the models alongside concurrent daily administration of GLYB, an inhibitor of the SUR1-TRPM4 channel. This channel is thought to have protective effects against chronic seizure development (*17-19*). We selected GLYB treatment for this comparison, rather than using a standard antiseizure medication, which would affect standard PTZ kindling, because preliminary experiments showed no response to GLYB treatment in the standard model, unlike its effects on other epilepsy models (*17*). Therefore, we aimed to determine whether the PTZ-DE model could detect antiseizure effects of GLYB treatment on chronic seizure susceptibility that were missed by the standard approach. In the standard PTZ kindling setup, concurrent GLYB treatment did not alter kindling rates (p=0.919, F=0.1671) (Figure 3A). However, with the PTZ-DE model, mice co-treated with GLYB exhibited a significantly reduced seizure response (p<0.001, F=15.29), characterized by a rightward shift in the kindling response curve and decreased Racine scores throughout, including the maximal plateau (Ymax: VEH: 4.527; GLYB: 4.025, p=0.03, F=4.927, extra sum-of-squares F-test) (Supplemental Table 1). Additionally, GLYB-treated mice undergoing the PTZ-DE model required significantly more 30mg/kg PTZ administrations to achieve Racine 4 or above seizures (p=0.04) compared to vehicle treated mice (Figure 3C). As with the experiments in section 3.1, PTZ-DE kindling curves demonstrated a better fit with the logistic growth model (R^2^: VEH: 0.7957; GLYB: 0.7404) compared to standard kindling curves (R^2^: VEH: 0.3334; GLYB: 0.2792). These findings indicate that the PTZ-DE model offers enhanced sensitivity to detect subtle chronic seizure threshold changes in response to pharmacologic treatment compared to the traditional PTZ kindling approach.

**Figure 3:**
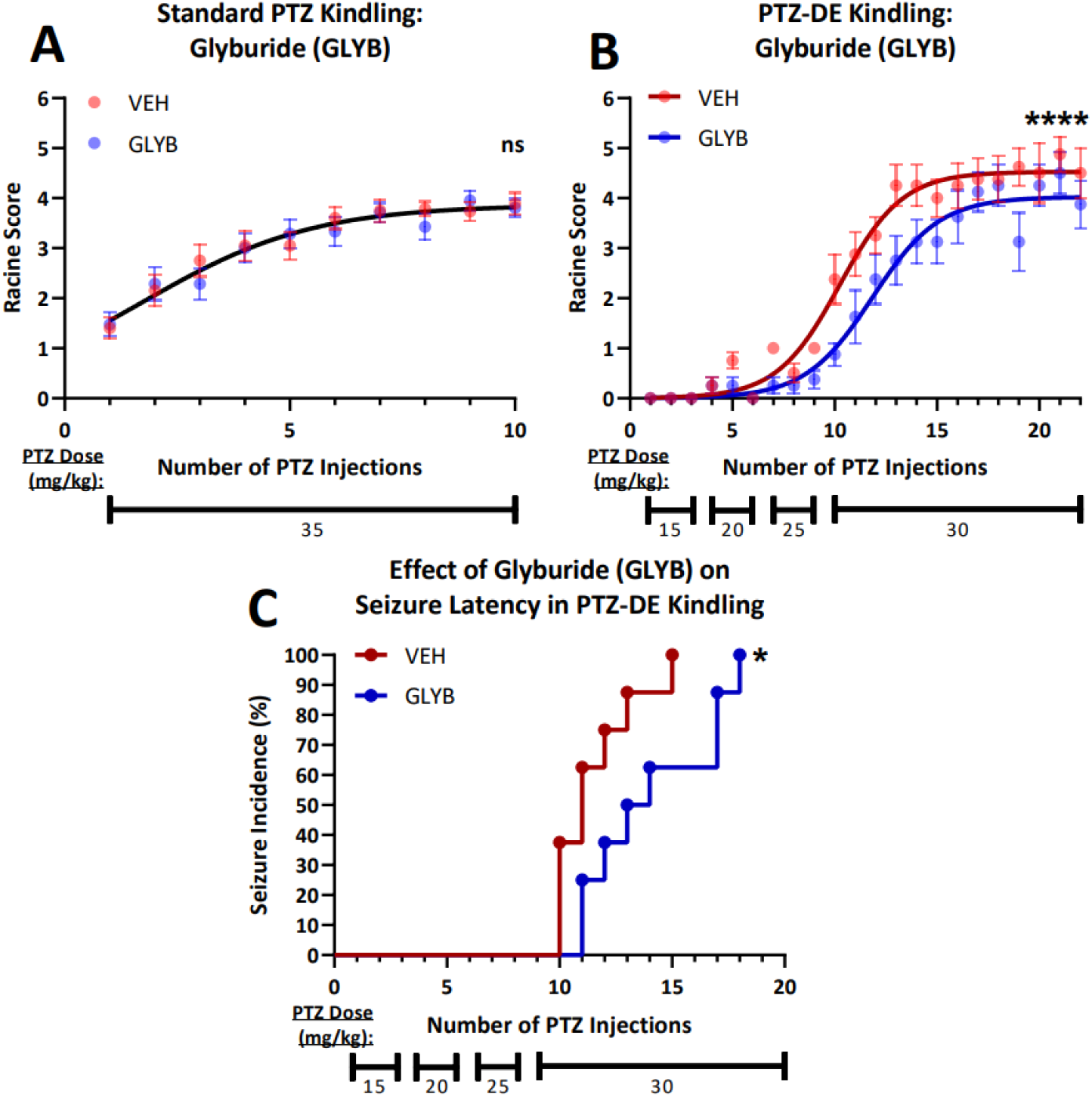
The PTZ-DE Model Improves the Detection Sensitivity of PTZ Kindling to Detect Underlying Differences in Chronic Seizure Susceptibility. A) Mice undergoing concurrent daily glyburide (GLYB) administration (n=20) demonstrate similar seizure response to standard PTZ kindling compared to vehicle controls (n=20). B) Mice undergoing concurrent daily GLYB administration (n=8) with PTZ-DE kindling demonstrate significantly less severe seizure response compared to vehicle control (n=8) mice. C) Daily GLYB treatment increases the number of PTZ administration in the PTZ-DE model required to induce the first Racine 4+ (tonic-clonic) seizure response. ns: not significant *p<0.05, ****p<0.0001.

## 4. Discussion

The PTZ kindling method serves as a reliable model for assessing chronic seizure susceptibility, independent of status epilepticus. In this study, we introduce a novel PTZ-DE model which employs progressively increasing doses of PTZ to empirically identify the lowest convulsive dose required for kindling under varying experimental conditions (Figure 1). This approach not only replicates the effects observed with the standard PTZ kindling model (Figure 2) but also enhances the detection of differential responses across diverse applications (Figure 3). Consequently, the PTZ-DE model may offer enhanced translational potential by improving the ability to identify nuanced differences in genetic or pharmacological therapies, thereby uncovering novel targets and minimizing the risk of missing opportunities for preventing or treating epilepsy. The PTZ-DE model improves on the traditional PTZ kindling approach by offering greater sensitivity and applicability across a broader range of animals and experimental conditions. Variability in group responses to PTZ, influenced by factors such as age, sex, and genetic background, poses challenges in the standard PTZ kindling model, which assumes uniform responses to a fixed PTZ dose (*4-7*). This assumption can introduce variability between groups when using animals with diverse characteristics, such as genetic knockout models (Figure 1). By tailoring PTZ doses to the specific attributes of animals in each experiment, the PTZ-DE model allows for the detection of subtle differences in chronic seizure susceptibility (Figure 1, 3). Additionally, the PTZ-DE model improves the sensitivity by increasing the total number of PTZ administrations, thereby enhancing degrees of freedom and statistical power (Supplemental Table 1). Importantly, by correcting for variations in seizure responses caused by diverse genetic backgrounds, the PTZ-DE model offers improvements in experiments using genetic knockout models bred on mixed backgrounds.

Beyond improving sensitivity, the PTZ-DE model also better replicates the expected kindling response compared to standard PTZ kindling, ideally represented by an S-shaped curve modeled through logistic growth regression (Figure 1, Supplemental Table 1). As demonstrated throughout our data, PTZ-DE curves consistently maintain a better fit to the observed data (represented by R^2^ and RMSE values) compared to standard PTZ curves (Supplemental Table 1). Starting with a lower PTZ dose followed by gradual escalation, the PTZ-DE model produces minimal seizure responses (Racine 0) at the beginning of kindling (represented in the logistic growth model by a low Y_0_ value) until the minimal effective dose is reached (Figure 1, Supplemental Table 1). Once reached, seizure severity increases and eventually plateaus (Ymax) as the “fully kindled” state is reached (Figure 1, Supplemental Table 1). In contrast, the standard PTZ kindling model exhibits an elevated Y_0_ values due to the typically used starting dose of 35mg/kg, which is higher than the minimally effective dose in most cases (Figure 1, Supplemental Table 1). This elevated initial response limits the ability to model the chronic lowering of seizure thresholds—a key feature of epileptogenesis. The PTZ-DE model overcomes this issue by empirically determining the minimally effective dose, ensuring the starting dose for kindling is at or below that dose and accurately captures the phase of chronic seizure threshold reduction.

An alternative approach to empirically determining the optimal PTZ dose could be to conduct a series of pilot studies to establish a PTZ dose response curve, with either acute or chronic PTZ exposure, and using this to inform a standard PTZ kindling protocol (e.g., targeting an EC25 point). However, this approach would not account for cohort variability in PTZ response and would be time and cost prohibitive, especially in conditions where breeding is required for experiments. The PTZ-DE model is a superior approach because it determines the ideal PTZ dose for a given experiment not only within varied animal backgrounds, but also within the exact set of animals being used for a given experiment. This is cost-effective and eliminates the effect of cohort variability in response to PTZ.

Despite its strengths, a limitation of PTZ kindling, and thus the PTZ-DE model, is the lack of consistent spontaneous seizures, a hallmark of epilepsy in patients (*20*). Alternative chemical induction models such as lithium pilocarpine or kainic acid, which reliably produce spontaneous seizures are often used (*21*). However, these models require inducing status epilepticus and/or directly injecting the brain, complicating the distinction between chronic epilepsy mechanisms and those resulting from primary injuries like status epilepticus or trauma (*21*). These limitations narrow their translational applicability (*22*). In contrast, PTZ-DE kindling simulates chronic epileptogenesis without requiring a primary injury, making it broadly applicable to a variety of chronic epilepsies (*3, 21*).

The primary limitation of this study is the absence of electrographic characterization of the PTZ-DE model, which we intentionally avoided to prevent confounding effects from intracranial electrode implant (*23*). This omission means potential unique electrographic events or spontaneous seizures induced by this modified kindling approach may have been missed. Given the standard PTZ models known limitation in reliably producing spontaneous seizures, future studies should incorporate long-term electrographic monitoring of the PTZ-DE model. This would clarify whether the increased number of PTZ administrations elevates the model’s severity and potentially induces spontaneous seizure activity.

## 5. Conclusions and Future Directions

The PTZ-DE model is a novel method designed to evaluate changes in chronic seizure thresholds, building upon traditional PTZ kindling by incorporating dose escalation, a concept adapted from acute seizure threshold testing. This model enhances sensitivity in detecting differences in seizure susceptibility across various conditions and allows for customization of PTZ dosing tailored to specific experimental conditions. As a result, the PTZ-DE model has the potential to facilitate the discovery of new therapeutic targets for the treatment and prevention of epilepsy.

## Abbreviations

PTZ: Pentylenetetrazol
PTZ-DE: Pentylenetrazol Dose Escalation
IP: Intraperitoneal
TBI: Traumatic Brain Injury
GLYB: Glyburide
VEH: Vehicle
CCI: Controlled Cortical Impact
AICc: Akaike’s Information Criterion
GRC: Growth Rate Constant

## Acknowledgments

JMS is supported in part by a grant from the National Institute of Neurological Disorders and Stroke (R01NS127986). MM is supported in part by a grant from the National Institute of Neurological Disorders and Stroke (1F31NS135752-01A1).

## Author Contributions

Conceptualization, M.M., M.B., V.G., J.M.S., A.K.; formal analysis, M.M., C.C.; investigation, M.M., J.L., O.T., D.M.; data curation, M.M.; writing—original draft preparation, M.M., J.L., D.M.; writing—review and editing, M.M., J.M.S., A.K.; visualization, M.M.; funding acquisition, J.M.S., A.K. All authors have read and agreed to the published version of the manuscript.

