## Supplemental Table 1 for "Dose Escalation in Pentylenetetrazol Kindling Detects Differences in Chronic Seizure Susceptibility"

**Supplemental Table 1: Complete Statistical Description for All Comparisons in This Study**

| | | | | | | | | Effect Size | | | Variability/Fit Measure | |
|---|---|---|---|---|---|---|---|---|---|---|---|---|
| Figure | Comparison | Sample Size | Test Used | Conclusion/Interpretation | P-Value | F-Value | Degrees of Freedom | Growth Rate Constant (k) | $Y_0$ | Ymax | $R^2$ | Chi-Square |
| 1A | Genetic Strain (C57/Bl6N vs C57/Bl6N+SvJ) PTZ Standard | C57/Bl6N: 5<br>C57/Bl6N+129/SvJ: 5 | Logistic Growth, Least Square Fit | Reject the null hypothesis; Different curve for each data set | <0.0001 | 8.964 | C57/Bl6N: 47<br>C57/Bl6N+129/SvJ: 47 | C57/Bl6N: 0.5319<br>C57/Bl6N+129/SvJ: 0.6684 | C57/Bl6N: 0.6160<br>C57/Bl6N+129/SvJ: 2.076 | C57/Bl6N: 4.835<br>C57/Bl6N+129/SvJ: 4.152 | C57/Bl6N: 0.5826<br>C57/Bl6N+129/SvJ: 0.1744 | N/A |
| 1B | Genetic Strain (C57/Bl6N vs C57/Bl6N+SvJ) PTZ-DE | C57/Bl6N: 14<br>C57/Bl6N+129/SvJ: 5 | Logistic Growth, Least Square Fit | Reject the null hypothesis; Different curve for each data set | <0.0001 | 14.47 | C57/Bl6N: 305<br>C57/Bl6N+129/SvJ: 97 | C57/Bl6N: 0.7196<br>C57/Bl6N+129/SvJ: 0.3419 | C57/Bl6N: 0.002903<br>C57/Bl6N+129/SvJ: 0.1681 | C57/Bl6N: 4.409<br>C57/Bl6N+129/SvJ: 3.814 | C57/Bl6N: 0.8262<br>C57/Bl6N+129/SvJ: 0.6366 | N/A |
| 1C | Sex (M vs F) PTZ Standard | Male: 5<br>Female: 8 | Logistic Growth, Least Square Fit | Reject the null hypothesis; Different curve for each data set | <0.0001 | 10.86 | Male: 57 Female: 77 | Male: 0.08362<br>Female: 0.3765 | Male: 1.374 Female: 1.744 | Male: 3.717E+86 (unstable)<br>Female: 4.368 | Male: 0.1670<br>Female: 0.2508 | N/A |
| 1D | Sex (M vs F) PTZ-DE | Male: 7<br>Female: 7 | Logistic Growth, Least Square Fit | Do not reject the null hypothesis; One curve for all data sets | 0.2350 | 1.427 | Male: 151<br>Female: 151 | Male: 0.7096 Female: 0.7724 | Male: 0.002685<br>Female: 0.002089 | Male: 4.544 Female: 4.265 | Male: 0.8158<br>Female: 0.8436 | N/A |
| 2A | SHAM vs TBI PTZ-DE Racine | SHAM: 8<br>TBI: 8 | Logistic Growth, Least Square Fit | Reject the null hypothesis; Different curve for each data set | <0.0001 | 16.29 | SHAM: 157 TBI: 157 | SHAM: 0.1462 TBI: 0.5483 | SHAM: 0.6277 TBI: 0.04763 | SHAM: 7.265 TBI: 5.253 | SHAM: 0.4382 TBI: 0.5979 | N/A |
| 2B | SHAM vs TBI PTZ-DE Mortality | SHAM: 8<br>TBI: 8 | Log-rank (Mantel-Cox) test | Reject the null hypothesis | 0.0297 | N/A | N/A | N/A | N/A | N/A | N/A | 4.726 |
| 3A | VEH vs GLYB PTZ Standard | VEH: 20<br>GLYB: 20 | Logistic Growth, Least Square Fit | Do not reject the null hypothesis; One curve for all data sets | 0.9185 | 0.1671 | VEH: 194<br>GLYB: 207 | VEH: 0.5739 GLYB: 0.4901 | VEH: 1.037 GLYB: 1.150 | VEH: 3.861 GLYB: 3.874 | VEH: 0.3334 GLYB: 0.2792 | N/A |
| 3B | VEH vs GLYB PTZ-DE Racine | VEH: 8<br>GLYB: 8 | Logistic Growth, Least Square Fit | Reject the null hypothesis; Different curve for each data set | <0.0001 | 15.29 | VEH: 173<br>GLYB: 173 | VEH: 0.6525 GLYB: 0.6089 | VEH: 0.005675<br>GLYB: 0.002982 | VEH: 4.527 GLYB: 4.025 | VEH: 0.7957 GLYB: 0.7404 | N/A |
| 3C | VEH vs GLYB PTZ-DE Seizure Latency | VEH: 8<br>GLYB: 8 | Log-rank (Mantel-Cox) test | Reject the null hypothesis | 0.036 | N/A | N/A | N/A | N/A | N/A | N/A | 4.398 |
